# Spatial proteomics of the human atherosclerotic microenvironment reveals heterogeneity in intra-plaque proteomes and extracellular matrix remodeling

**DOI:** 10.1101/2025.06.13.659602

**Authors:** Kathrine J Jokumsen, Lasse G Lorentzen, Karin Yeung, Timothy A. Resch, Jonas P Eiberg, Michael J Davies, Luke F Gamon

**Affiliations:** Department of Biomedical Sciences, Panum Institute, University of Copenhagen, Denmark; Department of Vascular Surgery, Heart Centre, Copenhagen University Hospital - Rigshospitalet, Copenhagen, Denmark; Department of Clinical Medicine, Faculty of Health and Medical Sciences, University of Copenhagen, Denmark; Copenhagen Academy for Medical Education and Simulation (CAMES), Capital Region of Denmark, Copenhagen, Denmark

**Keywords:** Spatial proteomics, laser capture microdissection, atherosclerosis, cardiovascular disease, proteomics, mass spectrometry, extracellular matrix, carotid endarterectomy

## Abstract

The heterogeneity of atherosclerotic plaques is critical for their vulnerability to rupture and the associate risk of cardiovascular events. Most proteomic studies have only examined *bulk* changes, potentially obscuring key spatial differences in protein content and abundance. Here we report a high-resolution spatial proteomics workflow that allows exploration of the molecular landscape of human plaques and murine myocardial tissue. This combines laser capture microdissection of tissue areas (50,000 µm² from 10 µm-thick sections, corresponding to < 30 cells), with high-sensitivity ion-mobility mass spectrometry, allowing spatial profiling of cellular and extracellular matrix (ECM) proteomes. Over 2700 proteins were detected, revealing substantial intra-plaque proteome heterogeneity across distinct regions (lipid-rich, media layers, shoulder regions, necrotic core, intima) and distance from the lumen into the artery wall. Strong inverse correlations were detected between proteases (e.g. cathepsin-B) and core structural ECM components (e.g. perlecan, HSPG2) consistent with active ECM remodeling. Analysis of media layers indicated distinct protein signatures associated with smooth muscle contraction and cell-cell communication. Blood coagulation signatures, including platelet degranulation and fibrin clot formation were enriched at the intimal surface. Inflammatory markers (clusters of differentiation 4 and 68, CD4/CD68; vascular cell adhesion molecular 1, VCAM1) and vascular damage markers (tenascin-C, TNC) were enriched in shoulder regions. The necrotic core was dominated by blood proteins, consistent with intra-plaque hemorrhage. The capacity of this workflow to resolve changes over modest distances (225 µm) provides unprecedented insights into the spatial organisation of the atherosclerotic microenvironment, offering a powerful tool for elucidating plaque biology and identifying potential therapeutic targets.

## Introduction

Cardiovascular diseases (CVD) remain the leading cause of death worldwide and a major global health challenge [1]. Atherosclerosis is the primary cause of cardiovascular disease which remains the leading cause of death worldwide and thus a major global health challenge [1, 2]. CVD has a complex pathophysiology and is characterised by the formation of arterial plaques with different complexity that can lead to plaque rupture, arterial thrombosis, peripheral embolisation and organ ischaemia [3, 4].

Plaques are highly heterogeneous with key features including lipid/cholesterol-rich fatty deposits, a necrotic core of debris, calcifications, intra-plaque hemorrhage, incorporated thrombi and the formation of a fibrous cap [3, 4]. Plaque rupture is often associated with ulceration, a thin fibrous cap, intra-plaque hemorrhage, a large necrotic or lipid core, low numbers of synthetic smooth muscle cells and large numbers of activated immune cells. In contrast, calcified or ‘hard’ plaques are typically regarded as more stable, and less prone to rupture. However, the molecular patterns underlying different plaque features and their association with plaque rupture remain to be explored in detail [3–5].

The early stages of plaque formation are characterised by endothelial dysfunction, accumulation of low-density lipoproteins (LDL) and formation of lipid-loaded (foam) cells, phenotypic alteration, migration and proliferation of smooth muscle cells, and immune cell infiltration and activation. The formation of a chronic inflammatory environment promotes the continued recruitment of activated immune cells, oxidant formation, and expression, secretion and activation of proteases [4, 5]. This results in remodeling and significant changes to the proteins, glycoproteins and proteoglycans of the extracellular matrix (ECM) that surrounds cells, and forms the fibrous cap. An imbalance between the rate of synthesis and degradation of the ECM of the fibrous cap towards the latter can decrease the stability of the fibrous cap and enhance its propensity to undergo ulceration and rupture [6]. In contrast, greater rates of ECM synthesis and accompanying mineralisation can result in thick, calcified and stable fibrous caps that less likely to undergo rupture [5].

Gross plaque morphological features are important in clinical diagnosis, with ultrasound derived calcification scores used to estimate plaque burden in patients [7]. While the risk of rupture depends greatly on plaque composition, nearly all previous transcriptome and proteome studies have relied primarily on *bulk* analysis where entire plaques are homogenised prior to biochemical analysis [5]. Previous proteomic analyses of complete plaques, using liquid chromatography-mass spectrometry (LC-MS/MS), have provided evidence that ‘hard’ plaques show higher levels of structural ECM proteins, and those associated with calcification, whereas ‘soft/rupture-prone’ plaques contain higher levels of inflammatory markers, oxidant-generating species and proteases [8–12]. Furthermore, protein ‘degradomic’ analyses (the complement of fragments from protein turnover/degradation) have provided evidence for characteristic and elevated levels of protein fragments from ECM and cellular proteins in ‘soft/rupture-prone’ plaques [13].

Recent work has shown distinct gene expression profiles in rupture-prone carotid artery plaques proximal to, or at the site of greatest narrowing, when compared to more stable distal areas, highlighting the critical role of *localised* molecular process in plaque vulnerability [14]. Integrated analysis of single-cell and spatial transcriptomics have revealed the importance of the microvasculature and its interaction with immune and smooth muscle cells in plaque development [15, 16]. While these transcriptomics approaches offer valuable insights into the importance of plaque microenvironments, they offer an incomplete picture especially with regards to ECM molecules where there can be marked discrepancies between gene and protein expression, and also for proteases, where RNA data provides little data on activity [17, 18]. Few studies have provided localised proteome data for plaques, though it has been reported that peripheral regions of carotid artery plaques (distal or proximal ends) differ in their proteome compared to the core (middle or most occluded part of the plaque) [19].

Here, we outline a workflow and proof-of-concept for LC-MS/MS based spatial proteomics (protein mapping) of murine myocardial tissue and human carotid plaques by combining laser-capture microdissection, limited sample handing and high-sensitivity ion-mobility mass spectrometry.

## Methods

### Ethics

The human studies were conducted according to the Helsinki declaration, and data and biological materials were collected after patients’ written, verbal, and informed consent. Study approval was obtained from The Danish National Committee on Health Research Ethics (journal number H-20002776). The animal studies were approved by the Animal Experiments Inspectorate, Ministry of Food, Agriculture and Fisheries, Denmark (approval number 2021-15-0201-00818).

### Tissue collection, embedding and sectioning

During carotid endarterectomy (CEA) for symptomatic ipsilateral internal carotid artery (ICA) stenosis, the atherosclerotic plaque was removed in toto and then processed as described in the Supplementary Methods. Patients were recently diagnosed with cerebral embolic ischaemia (stroke, transient ischaemic attack) or retinal ischaemia (complete vision loss, amaurosis fugax). Diagnosis and medical therapy were carried out by one of four referring neurological departments, but all surgical procedures were carried out by the same department of vascular surgery as described recently [8]; for further details see Supplementary Data.

B6.129P2-*Apoe^tm1Unc^* N11 male mice aged 8 weeks were purchased from Taconic Biosciences. Animals were housed in a temperature-controlled facility with a 12h light/dark cycle and received food and water *ad libitum*. The mice were euthanised at 24 weeks, perfused with cold saline before the heart was collected and processed as described in the Supplementary Methods.

### Laser capture microdissection

Laser capture microdissection and microscopy was performed with a Zeiss PalmRobo and 5x objective. Individual areas were cut and loaded into water droplets in 8-well flat-cap PCR strips. Matched tissue areas were collected from up to 7 sequential sections. For further details, see Supplementary Methods.

### Proteomic sample preparation

Initial studies employed small tissue sections generated by laser-capture microdissection (LCM) of formaldehyde-fixed 10 μm thick slices from murine myocardium. Circular areas of increasing size were excised, collected individually into water droplets in 8-well PCR-cap stripss. Samples were processed using a ‘hanging-drop’ method in a humid chamber [20], with the samples treated with 100 mM TEAB and 0.05% DDM (70°C, 40 min), followed by reduction and alkylation (10 mM TCEP, 40 mM CAA, 70°C for 20 min), digestion with trypsin (overnight, 37°C), followed by acidification with 10% TFA. This hanging drop method circumvented problems in the handling and centrifugation of small tissue samples and facilitated sample clean-up in 96-well format (see Supporting Data), which was performed by centrifuging directly onto home-made stage-tips in a 96-well format, minimising sample transfers and potential losses [21, 22]. Samples were subsequently reconstituted in 5% formic acid in water. For further details, see Supplementary Methods. This approach is similar in concept to published One-Tip and Chip-Tip methods [23, 24] but uses readily-available and low cost labware, when compared to these EvoSep-based separation methods. The volume of the captured areas from the murine heart sections is equivalent to 30 - 40 cardiomyocytes calculated using a mean volume of 13100 - 16800 μm^3^ for these cells[25].

One fifth of the sample was subsequently analysed. Thus the data reported correspond to the protein complement of 6-8 cardiomyocytes for the smallest sample sections analysed. For smaller cell types (e.g. HeLa cells), the corresponding number is approximately 20 cells. Protein extraction yielded 4 - 30 ng of protein depending on the size of the captured area.

### Liquid-chromatography mass spectrometry proteomic analysis

Samples were analysed on a TimsTOF Pro mass spectrometer (Bruker Daltonics) in the positive ion mode after separation using a C18 column at room temperature with gradient elution using 0.1% formic acid in water (Solvent A) and 99.9% acetonitrile/0.1% formic acid (Solvent B). Data were obtained using either data-dependent acquisition with parallel acquisition-serial fragmentation (DDA-PASEF) or data-independent acquisition parallel acquisition-serial fragmentation (DIA-PASEF) modes [26, 27]. DDA-acquired data were searched against the human UniProt reference proteome including common contaminants with a false discovery rate (FDR) set at 1% (protein and peptide level), and both fixed and variable modifications (see Supplementary Methods) [28–33]. DIA data were searched against the human UniProt reference proteome using DIA-NN version 1.9.1 (see Supplementary Methods). In accordance with Danish legislation relating to the Guidelines for Data Protection Regulation (GDPR), raw mass spectrometry data can only be shared under a data processing agreement between the data controller and processor. Such data sharing agreements are available and can be completed by contacting the corresponding authors.

### Statistical analysis and visualisation

Data analysis and visualisation of DIA data were performed using R (version 4.4.0) and R-Studio (Mac version 2024.04.1+748) to generate lists of differentially expressed proteins using a robust linear model approach. Proteins were considered as expressed differentially if the Benjamini-Hochberg adjusted p-value was < 0.01 [34]. Further details are provided in the Supplementary Methods. Gene Set Enrichment Analysis (GSEA) was carried out using the clusterProfiler/ReactomePA packages using median-normalised log2 fold changes as input, and adjusted p-values <0.05 as cut-off.

## Results

### Development of a high-throughput spatial proteomics workflow using murine myocardial sections

The workflow used for method development is outlined in Figs. 1 and 2. Mass spectral data, obtained in data-dependent acquisition (DDA) mode, allowed median identification of 648 proteins from the smallest areas examined (50.000 µm^2^ of 10 µm thickness cut from murine myocardial tissue sections using laser capture microdissection), with this increasing to 1284 proteins for 200.000 µm^2^ (Fig. 2C). A strong overlap of protein identifications was observed, with most identified proteins present as a subset of the largest area (200.000 µm^2^) and only 40 identifications unique to the smaller tissue areas (50.000 or 100.000 µm^2^). A significant increase in protein identifications from 648 to 1047 proteins (for the smallest area of 50.000 µm^2^) was achieved by acquiring mass spectra using a data-independent acquisition mode, with parallel acquisition and sequential fragmentation (DIA-PASEF) (see Supporting Data).

**Figure 1.**
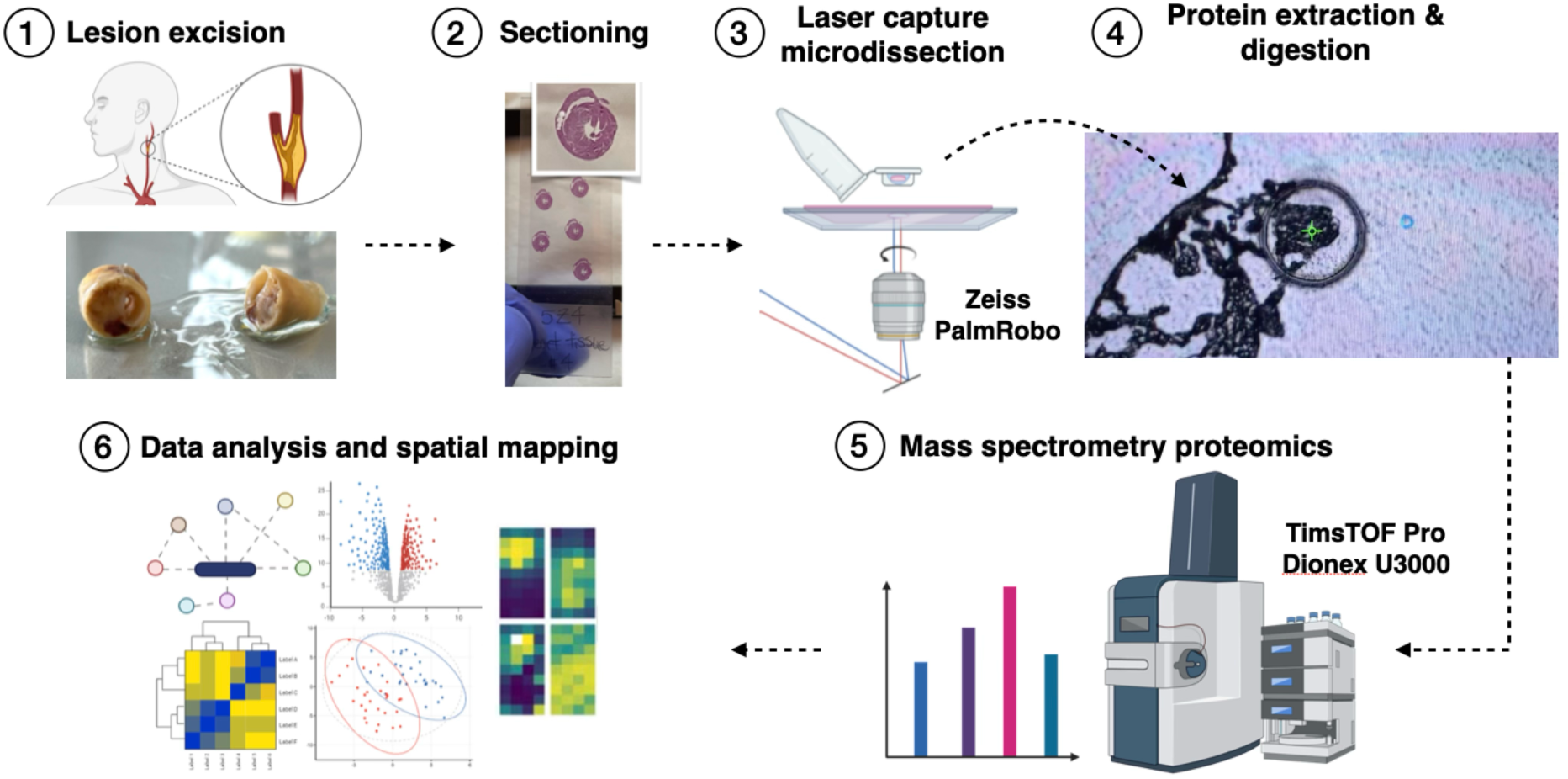
General workflow and overview. (1) Human carotid artery atherosclerotic plaques are excised from patients, (2) fixed and cryosectioned at 10 µm thickness. (3) Laser capture microdissection is used to excise 225 x 225 µm tissue areas (50000 µm^2^). (4) Proteins are extracted and digested as a hanging drop, followed by desalting in 96-well format. (5,6) Peptides are analyzed by liquid-chromatography mass-spectrometry, and the resulting data analysed.

**Figure 2.**
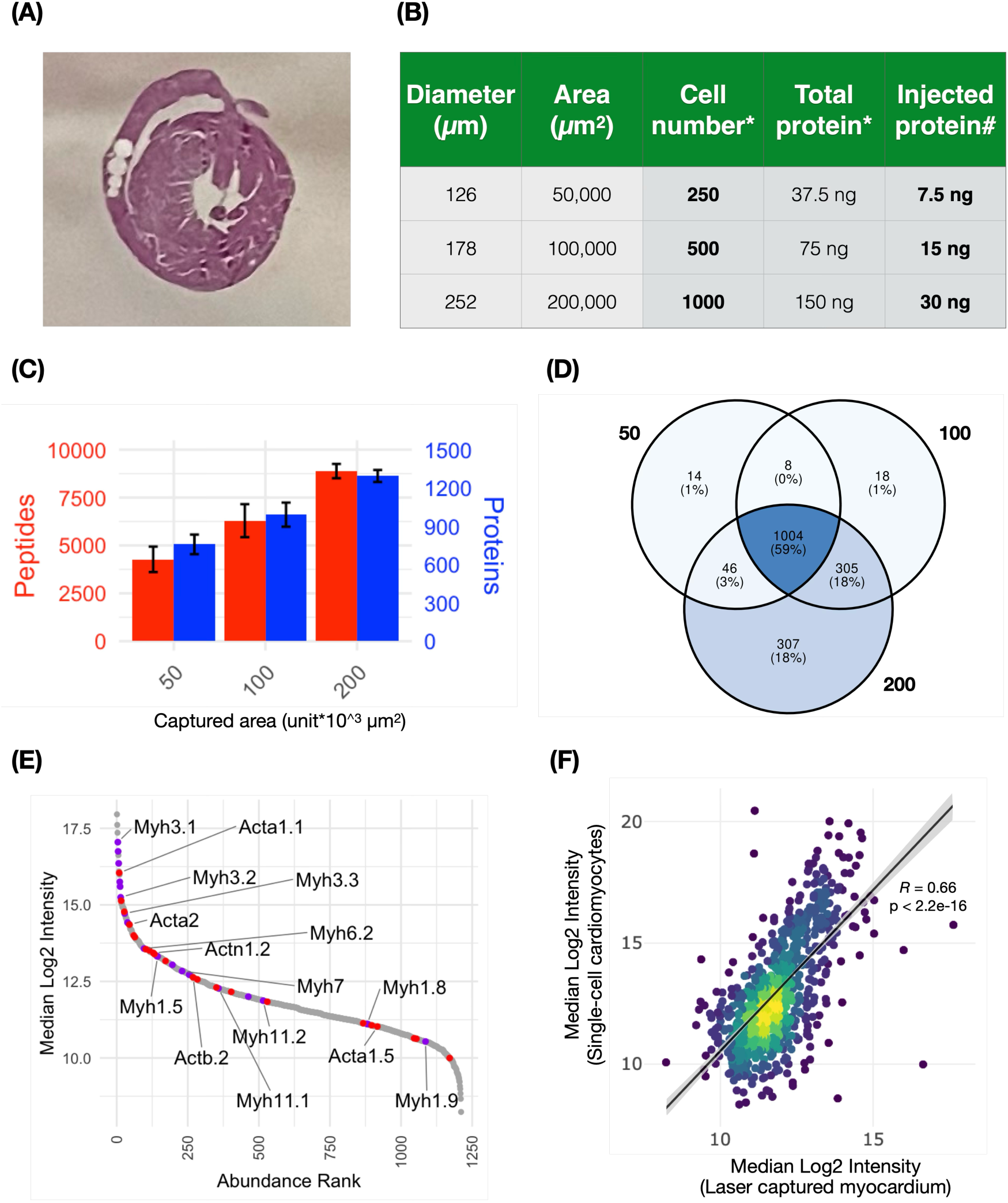
Optimisation of sample preparation and mass spectrometry parameters on formalin-fixed mouse heart sections. (A,B) Circular areas are removed by laser captured micro dissection from 10 µm thick sections of mouse heart. The captured regions of increasing areas of 50000, 100000, 200000 µm^2^ correspond to approximately 250-1000 cells, respectively, ∼7.5-30ng ng of protein. The estimated cell numbers are based on the tissue volume collected and average volume of a HeLa cell. One fifth of the sample was injected on-column. (C) Peptide and protein level identifications from the mouse myocardium areas. Increasing numbers of peptides and proteins were identified (using DDA-PASEF) with increasing area. (D) Venn diagram of protein identifications (using DDA-PASEF) from 50, 100 and 200 (*10^^3^ µm^2^) tissue areas show identifications are mostly subsets of those from larger samples. (E) The proteome of 50000 µm^2^ mouse myocardium tissue (DIA-PASEF) is dominated by myosin and actin muscle proteins. (F) The current data correlate strongly with that reported previously for single cell mouse cardiomyocytes (Kreimer et. al. Anal. Chem., **95**, 24, 9145-9150 (2023)).

Analysis of fixation artefacts (arising from formaldehyde, formylation or methylation; see Supplementary Data) indicated that these only affected a minor portion of the peptide spectral matches (PSMs). The proteome of the laser-captured mouse myocardium was dominated by actin and myosin species (Fig. 2E) and showed a strong positive correlation (*R* = 0.66; p < 2.2e-16) with published single-cell mouse cardiomyocyte proteomic data from Kreimer et al[35].

### Protein distributions at the lumen-plaque interface in human carotid atherosclerotic plaques

The above studies were subsequently extended to a more detailed proof-of-concept of protein mapping using symptomatic human carotid atherosclerotic plaques, with protein distributions examined at the lumen-intima interface and with increasing depth into the plaque. Tissue was collected in 2×4 square arrays of 50.000 µm^2^ area (Fig. 3). Subsequent LC-MS/MS analysis indicated a high degree of similarity between immediately adjacent samples collected at the same distance from the vessel lumen. Principal component analysis (PCA) and hierarchical clustering showed good clustering of differentially abundant proteins, and with distance from the lumen (see Supporting Data). This suggests that in this microenvironment, distance to the lumen is the major driver of proteome differences, likely reflecting changes in cell population from endothelial cells at the lumen interface, and lipid-rich (foam) cells in plaque, with corresponding enrichment of endothelial and adhesion-associated proteins at the intimal surface, and lipoproteins and collagens deeper into the plaque (Fig. 3B,C).

**Figure 3.**
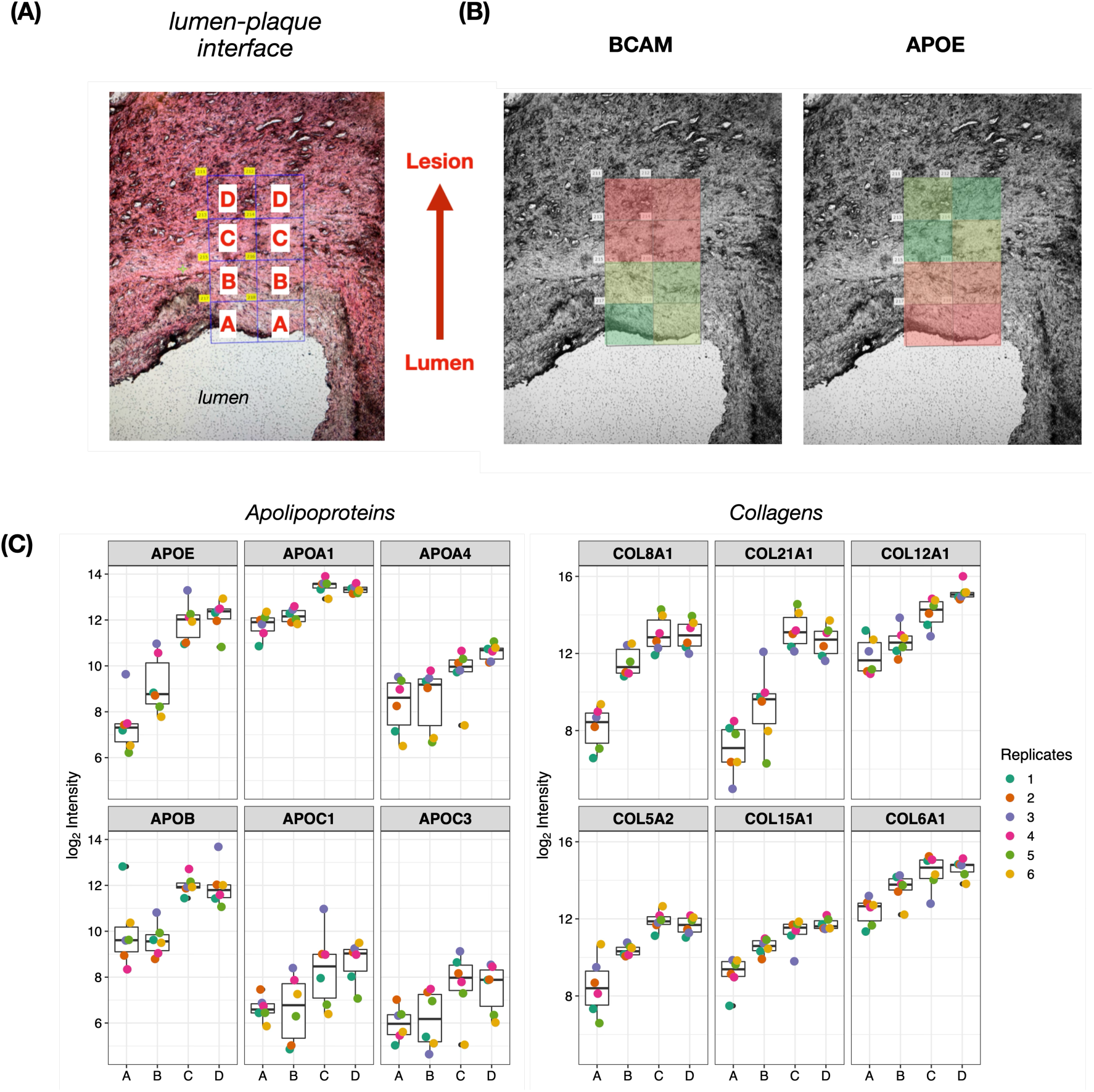
Spatial distribution of lipoproteins and collagens at the intima-plaque interface of a human carotid artery plaque with distinct abundance gradients on progressing from the lumen into the tissue. (A) 4×2 arrays were collected by LCM from sequential sections and samples were grouped based on distance from the lumen. (B) Representative spatial maps (Green = high intensity, Red = low intensity, log2Intensity scaled by protein); Basal Cell Adhesion Molecule (BCAM) is enriched at the luminal surface whereas Apoliprotein-E (APOE) is enriched deeper into the plaque. (C) Apolipoproteins (APOE, APOA1, APOA4, APOB, APOC1, APOC3) and collagens (COL8A1, COL21A1, COL12A1, COL5A2, COL15A1, COL6A1) are enriched within the plaque and increase in abundance with increasing distance from the lumen.

### Spatial proteomics reveals intra-plaque proteomic heterogeneity

Subsequent analyses examined various sites around the circumference of artery cross-sections (i.e. plaque, opposite wall, and intervening shoulder regions), with five morphologically-distinct regions examined, using 1×4 or 2×2 square arrays of 50.000 µm^2^ area (Fig. 4A). These included: 1) media and endothelium (mediaSq, 2×2 array); 2) intact smooth muscle cell rich media (mediaLn, 1×4 array); 3) plaque shoulder (shoulder, 2×2 array); 4) lipid and foam cell rich region (lipid, 2×2 array); and 5) necrotic and hemorrhaged core (necrotic, 2×2 array). Samples were acquired from sequential arterial cross-sections, with these used to obtain pseudo-technical replicates. Matched regions were removed by LCM, based on the alignment of morphological features. More than 2700 unique proteins were detected across the groups, with good clustering of the data in PCA analyses *within* a region (Fig. 5A), but large variances in identifications *between* regions (Fig. 4C). The highest numbers of protein identifications were obtained for the shoulder regions (purple), followed by the two media sections (pale and dark green), with lower numbers for the lipid-rich (red) and necrotic core (blue). Analysis of overlapping identifications revealed a high degree of proteome heterogeneity across the plaque (Fig. 4E). Of the identified proteins, approximately 33% were identified in all groups, similar numbers in the media or shoulder regions, and the remainder distributed more broadly across groups. A full list of the proteins detected and their spatial distributions is provided in Supplementary Table 1. Key inflammatory markers including cluster of differentiation 4 (CD4), cluster of differentiation 68 (CD68) and vascular cell adhesion protein 1 (VCAM1) were identified exclusively in the shoulder regions. The multi-dimensional PCA analyses (Fig. 4B) showed greatest separation between the cell-rich (mediaLn, mediaSq, shoulder), less diseased /more normal regions and the lipid-rich/necrotic/cell-poor regions.

**Figure 4.**
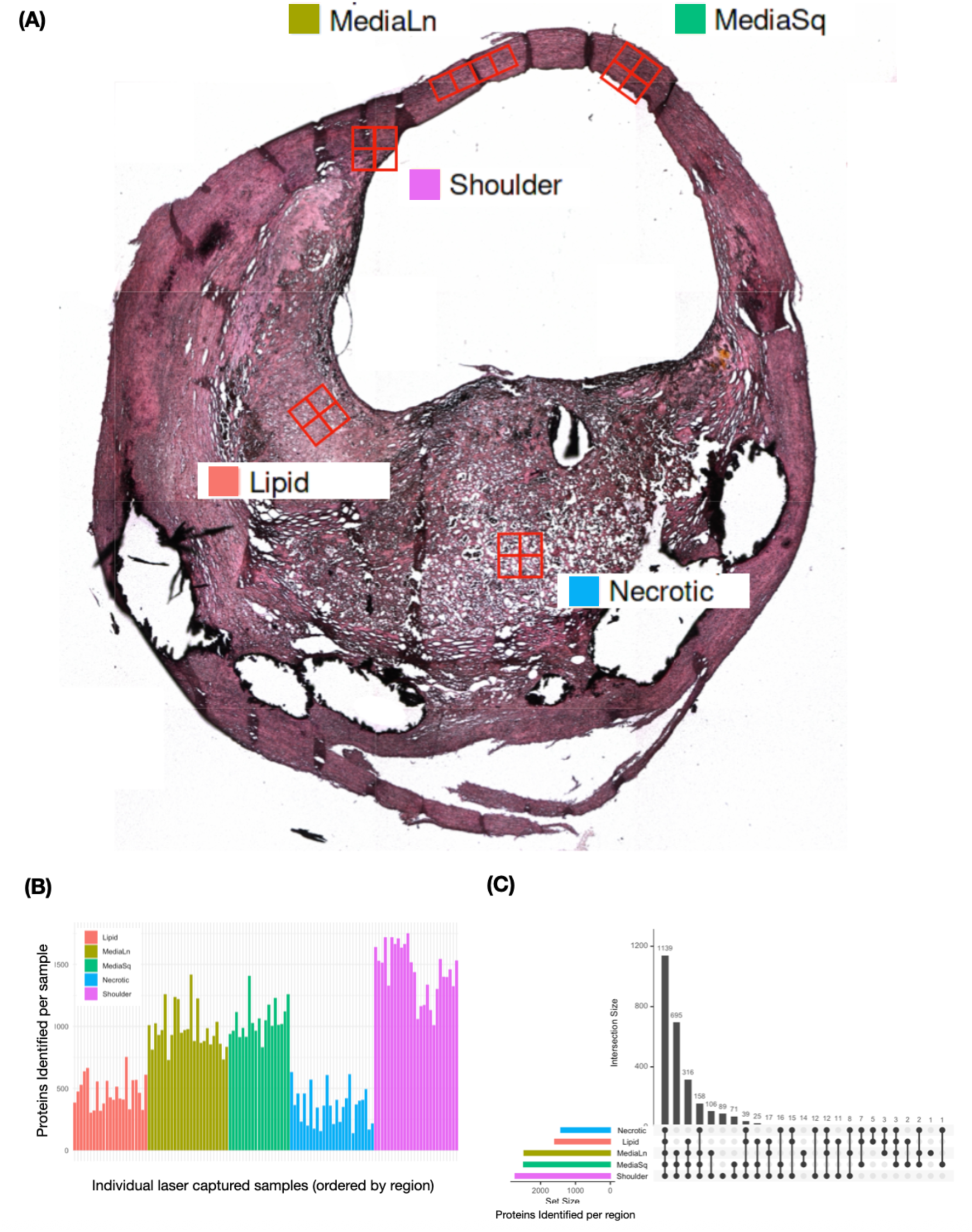
Spatial proteomics of a complex human carotid artery atherosclerotic plaque reveals protein heterogeneity. (A) Five distinct morphological areas were captured from H&E stained 10 µm thick tissue sections of a complex human plaque: **Lipid** (brick red color), foam cell and lipid rich region, **Necrotic** (blue color) necrotic core and intra-plaque hemorrhage, **Shoulder** (pink color) developing plaque shoulder region, **MediaLn** (olive green color) 4×1 linear array of side-by-side areas of intact media, **MediaSq** (bright green color) 2×2 square array containing two areas of media and two areas of endothelium. (B) Proteome coverage varies dramatically based on captured areas from ∼500-1500 proteins identified per individual sample. (C) UpSet plot of identified proteins grouped by region, showing majority of proteins identified in all regions or in the media associated areas (**MediaLn**, **MediaSq**, **Shoulder**).

**Figure 5.**
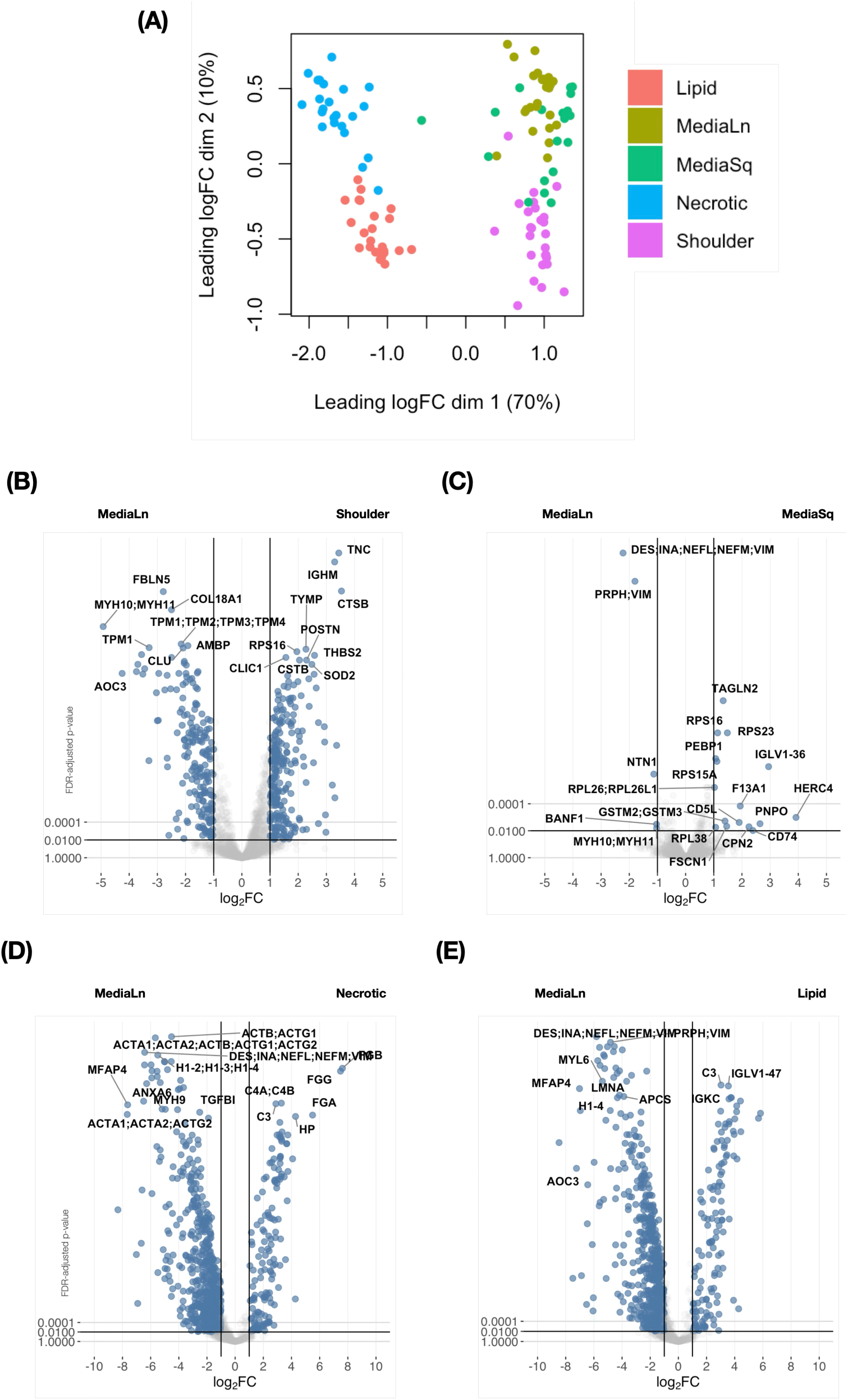
Principal component analysis and Volcano plots indicating differential protein expression compared to the intact media samples. (A) Areas separate according to morphological region by prinicipal component analysis (PCA) with clustering of media and shoulder areas. (B)-(E) Plots provide comparisons between the **Lipid**, **Shoulder**, **MediaSq** and **Necrotic** regions to the intact **MediaLn** region. Additional comparisons can be found in the Supporting Information.

The protein content in the plaque sections was compared to the more intact (‘normal/less diseased’) mediaLn region (Fig. 5). As expected, the related mediaSq region showed few statistically-significant differences in protein content (i.e. differentially-expressed species), indicating a close similarity between these samples (Fig. 5C). A total of 1053 proteins were differentially abundant across all comparisons, with the largest number of differentially abundant proteins been detected on comparison of the mediaLn against the lipid and necrotic regions (Fig. 5D,E). Comparison with the shoulder regions revealed increased abundance in the shoulder of known markers of vascular damage (TNC) and inflammation (IGHM, CTSB). The necrotic region was dominated by plasma proteins and those from red blood cells (C3, C4A/B, HP, FGA, FGB, FGG), consistent with intra-plaque, with thisconfirmed by gross visual assessment of the plaque. The lipid region was also dominated by highly abundant plasma proteins (Supplementary Table 1), though significantly higher levels of ECM proteins (e.g. COL6A1, COL6A2, COL6A3, LUM, BGN, POSTN) were observed on comparison of the lipid and necrotic regions, consistent with lipoprotein binding to ECM materials such as BGN [36], and ECM degradation in the necrotic region [6, 13].

Sub-region analysis of the intima and media areas within the mediaSq regions revealed many differentially abundant proteins (Fig. 6), with many involved in smooth muscle contraction, cell-cell interaction and ECM structure and function. Reactome pathway analysis showed ‘muscle contraction’, ‘smooth muscle contraction’ and ‘cell-cell’ communication pathways as the most significantly enriched in the media areas (Fig. 6C), with ‘extracellular matrix organisation’ and ‘platelet degranulation’ enriched in the intimal layer and vascular wall (Fig. 6D). Smooth muscle cell proteins (Fig. 6B; CNN1, LAMB2) and endothelial cell markers (Fig. 6B; VWF), were detected as enriched in the media and intima, respectively. Interestingly, we observed an enrichment in the intima of pathways associated with blood coagulation (‘platelet degranulation’, ‘formation of fibrin clot’) potentially indicating a contamination of the intimal surface with thrombotic material from a plaque rupture, or molecular signatures reflecting endothelial dysfunction or damaged intima.

**Figure 6.**
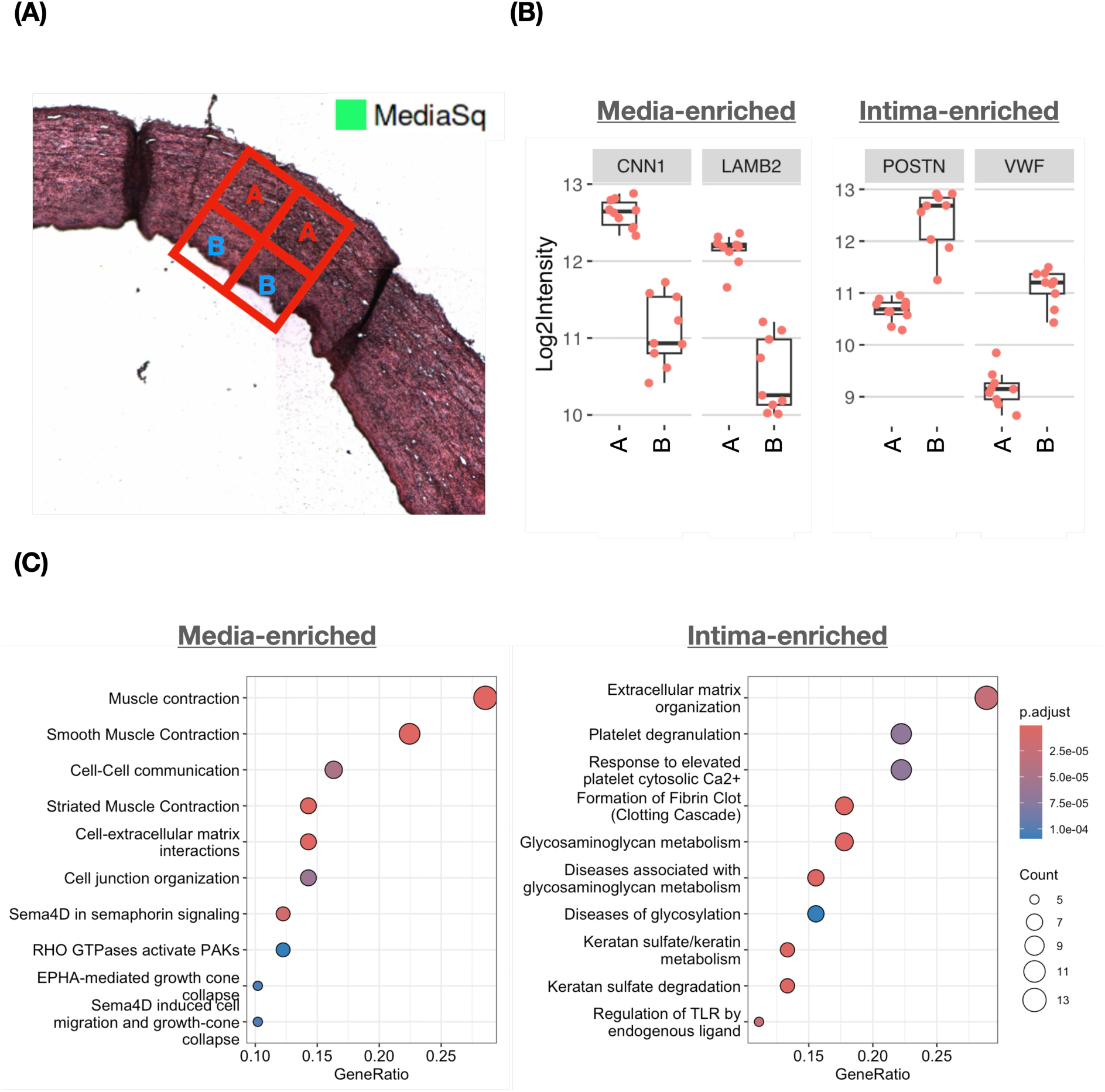
Spatial distribution of smooth muscle, extracellular matrix and blood proteins at the intimal-media interface. (A) Inset of a 2×2 array profiling the endothelium/intimal surface and smooth muscle rich media (**MediaSq**). (B) Example of smooth muscle cell proteins Calponin 1, Laminin Subunit Beta-2 (CNN1, LAMB2) enriched in the media locations and endothelial cell associated proteins Von Willebrand Factor, Periostin (VWF, POSTN) enriched in the endothelium. (C) Reactome pathway analysis performed on the statistically significantly different proteins show enrichment of muscle and cell interaction pathways in the media. Extracellular matrix, platelet degranulation and blood clotting pathways are enriched at the endothelium.

### Protein correlations reflect major cell populations and reveal immune-mediated extracellular matrix remodeling in the plaque shoulder

Large numbers of significant changes were observed in the proteomes detected for adjacent tissue sections in the shoulder region (Fig. 7). The greatest differences were observed between the developing shoulder (region ‘C’, Fig. 7A) and the area considered (on the basis of its location) to be most ‘media-like’ (region ‘B’, Fig. 7A). 334 proteins were differentially overabundant when comparing regions ‘B’ and ‘C’, with ‘B’ showing higher levels of proteins identified as being of smooth muscle cell origin (Supplementary Table 1). In contrast, markers of inflammation (MARCO, MMP12, CD36 and TNC) were more abundant in the developing shoulder areas ‘A’ and ‘C’, while MYH10 and VWF were enriched in areas of smooth muscle and endothelium, respectively (Fig. 7C).

**Figure 7.**
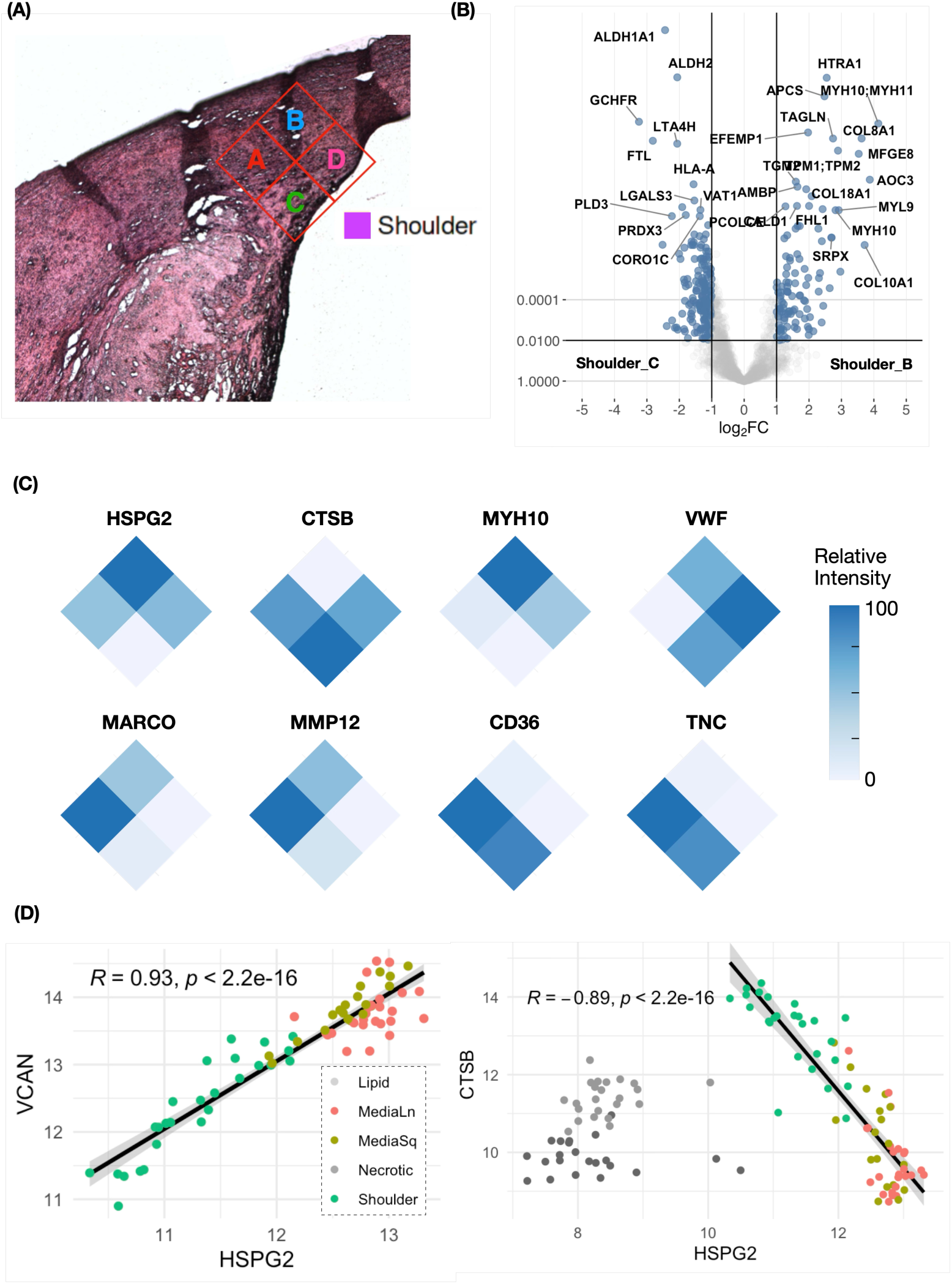
Inflammation and extracellular matrix remodeling changes in the shoulder. (A) Inset of shoulder and dissected areas. (B) Volcano plot of statistically significantly different proteins comparing location ‘B’ (media-like) and ‘C’ (shoulder/endothelium). (C) Spatial heat maps of selected proteins in the developing shoulder region (log2intensity values scaled to 0-100 by protein). Macrophage markers Macrophage Receptor With Collagenous Structure, Matrix Metalloproteinase-12 (MARCO, MMP12) are co-localized in the internal shoulder area, while smooth muscle and endothelial cell markers Myosin Heavy Chain 10, Von Willebrand Factor (MYH10, VWF) localize to endothelium and media-like areas, respectively. Markers of vascular dysfunction Cluster of Differentiation 36, Tenascin-C (CD36, TNC) are enriched in the the shoulder areas. (D) Scatter plots of Perlecan (HSPG2), an essential proteoglycan of the basement membrane, strongly positively correlating with Versican (VCAN) in media and shoulder areas, while Cathepsin-B (CTSB) inversely correlates is strongly inversely correlated. Scales are log2intensity.

Protein correlation analyses were performed across all samples, to identify spatially co-localised proteins. Global analysis was complicated by proteome heterogeneity and a high-degree of missing values in the necrotic and lipid regions (20 pairs with R^2^ > 0.95 when including all regions, 84 pairs with R^2^ > 0.95 when only including mediaLn, mediaSq and shoulder regions; and 121 vs 572 with R^2^ > 0.90, see Supporting Data). Amongst the most highly correlated protein groups in the mediaLn, mediaSq and shoulder regions were smooth muscle cell-derived proteins (e.g. MYL6, MYL9, MYH10, MYH11, FLNA, LAMC1 and LAMB2) presumably reflecting differences in smooth muscle cell density across these regions. HSPG2 and CTSB were highly *inversely* correlated when considering the mediaLn, mediaSq and shoulder regions, and the shoulder region alone (Fig. 7C,E), consistent with CTSB being involved in ECM degradation, and particularly of HSPG, whilst HSPG2 was highly correlated with other ECM proteins across the entire dataset (e.g. R^2^ > 0.97 with VCAN, Fig. 7D).

## Discussion

The sites of rupture of human atherosclerotic plaques are not uniformly distributed across the plaque surface area, with this occurring primarily at specific locations[14]. These include sites proximal to, and at, the site of maximum stenosis, but not distally (i.e. at locations after the site of maximum occlusion), and at the shoulders of the plaque where there is a transition from morphologically-normal arterial wall, to plaque[37]. This may be due to multiple factors including the extent of immune cell infiltration and activation, the extent of ECM remodeling or degradation, or as a result of high stress and turbulent blood flow arising from hemodynamic flow changes generated by the altered vessel morphology[38]. Despite this clear evidence for site-specific and localised events, many of the analyses carried out previously to examine plaque susceptibility, at both the transcriptome and proteome levels, have used *bulk* analyses, which may obscure key mechanistic details and pathways[15, 16].

The development of LCM has allowed the separation of specific regions for analyses, and this has been widely used to examine changes in transcriptome studies of plaques, however far fewer studies have used this approach, or others, in conjunction with LC-MS/MS to obtain a detailed (breadth and in-depth) analysis of the proteome of specific plaque regions[39, 40]. We report here a readily applicable workflow that provides proof-of-concept for spatial analysis of microenvironments within plaques, with these areas corresponding to low ng levels of proteins and <30 cells. The workflow was initially developed using fixed murine myocardial sections and subsequently extended to symptomatic human carotid artery plaques. In both cases, increasing the captured tissue volume enhanced the number of positive and robust proteins identifications, with even the smallest samples (approximately 125 μm diameter circular regions with an area of approximately 50000 μm^2^, yielding approximately 4 ng protein) allowing reproducible detection of approximately 600 proteins (Figs. 2B,C). The use of DIA-PASEF for acquiring MS spectra provided significant advantages over DDA, with an improvement in the detection of more than 1000 proteins from the mouse myocardium of the smallest area (50000 µm²). The strong correlation between laser-captured mouse myocardium, rich in cardiomyocytes, with single-cell proteome of sorted cardiomyocytes supports the robustness of this methodology. [35] The similarity is especially striking considering the analyses were performed in different labs, with different instruments and chromatographic methods, and mice with different genetic backgrounds.

The data obtained on examining sequential and adjacent sets of square LCM sections, generated from the lumen-endothelial interface and deeper into the center of human plaques, showed consistent proteome complements for parallel neighboring sections (both within single tissue slices, and for adjacent transverse slices) (Fig. 3). These showed the expected variation in proteins on progressing from the endothelial layer through the basement-membrane and into the lipid-rich regions with, for example, a decrease in the detection of basal cell adhesion molecule (BCAM) and increased levels of apolipoproteins and collagens (Fig. 3B,C).

Examination of morphologically-distinct regions within transverse artery/plaque sections, provided evidence of marked heterogeneity in both the number and types of proteins detected, with only approximately 33% of the total detected across all regions. The lowest numbers of protein identifications were for the lipid and necrotic core regions, possibly due to a lower concentration of intact proteins at these sites, or a reduced number of cell types (Fig. 4C). In contrast, the highest number of protein identifications were detected reproducibly in the shoulder regions, with this likely to be due to the diversity of cells present at such sites and also extensive ongoing inflammation. The sections obtained from the (less diseased, more normal) opposite artery wall (Fig. 4A) showed intermediate protein numbers, with these sections showing a higher degree of consistency in the identifications and a lower number of differentially expressed proteins (Fig. 5C). The UpSet plot in Fig. 4C illustrates the significant diversity of proteins identified across the regions, with this confirmed by the large number of differentially expressed proteins in the volcano plots (Fig. 5B,D,E). This heterogeneity indicates that bulk analyses, though giving important and useful data, may conceal critical differences in protein distributions.

A number of the differentially-expressed species have been associated previously with sites of vascular disease or vascular injury. Detailed analyses of 2×2 arrays dissected from the shoulder regions (Fig. 7) show even between these small sections (225 x 225 μm) there is a marked heterogeneity of protein species. Thus, HSPG2 and MYH10 showed the highest levels in region B (the most ‘media’ like) whereas VWF was highest in the endothelial-luminal interface region furthest from the plaque (region D). In contrast, regions A and C provided evidence for elevated levels of multiple inflammatory and tissue degradation markers including TNC which showed the greatest differential abundance in the shoulder regions (Fig. 5B) consistent with previous data [41]. The high abundance of TNC, possibly driven by Delta and Jagged signaling via the Notch receptor [42], may decrease cell adhesion and denudation via its known interactions with fibronectin [43, 44], and also enhance the expression of matrix metalloproteinases [45]. Consistent with this, these regions also contained high levels of inflammatory markers (e.g. CD36) and proteases (including CTSB and MMP12), and evidence for an increased extent of degradation of cell and ECM proteins (cf. the low levels of HSPG2). These data are consistent with the detection of large numbers of protein fragments in soft/vulnerable plaques in bulk analyses [13]. These shoulder areas also contained elevated levels of species associated with oxidation (e.g. GCFHR, which regulates nitric oxide production via regulation of tetrahydrobiopterin levels; PRDX3, which removes the oxidant H_2_O_2_), detoxification of reactive aldehydes possibly formed via lipid peroxidation (ALDH1A1, ALDH2), and the storage of damaging iron complexes (FTL). These differences are consistent with an increased susceptibility of the plaque to rupture in this region.

A number of the proteins identified in the ‘proteomic atlas’ dataset of Theofilatos et. al. [19] as differentially abundant in symptomatic plaque cores, were detected at a higher abundance in the shoulders when compared to the two ‘opposite wall’ media regions. Five of the twelve species were also differentially abundant in our dataset (CD14, CTSD, TNC, POSTN, CTSB) and an additional two were observed to have a fold change of > 0.2 and to be statistically significant with adjusted p-value < 0.05 (TIMP1, S100A8).

The current study, and the reported workflow, has a number of strengths and weaknesses. A strength of the workflow is the depth and reproducibility of the proteomic analyses from small tissue samples cut using LCM from fixed sections. As this type of tissue preparation is commonly used, and LCM equipment is relatively common, this protocol should be able to be readily adapted to many other tissue types. The capture and digestion protocol appear to be highly efficient and can be performed with standard labware, without the need for immunostaining or other specialised equipment such as that employed in the artificial intelligence/machine learning-enabled cell-centric laser capture methods employed, for example, in the Deep Visual Proteomics workflow [46].

The low amount of material required – both in terms of cell number and protein amounts (low ng) - are additional strengths. The resolution that is inherent in this method is a considerable strength, as it allows detailed mapping of protein changes across small distances (e.g. 225 μm; Fig. 7). The size of the sections that yield significant protein identifications is also a potential drawback, as complete mapping, even of single arterial cross-sections, generates enormous numbers of samples, which require considerable resources to process and analyse. A weakness of the current study, related to the above comment, is that it has only examined a small number of plaques and consequently plaque microenvironments within these, due to analytical limitations imposed by the numbers of samples generated. Furthermore, only symptomatic plaques have been examined, as asymptomatic subjects are not operated on in Denmark. Despite these limitations, this study has provided a clear ‘proof of concept’ for this approach, and rapid advances in laser capture automation and throughput, as well as LC-MS/MS instrumentation, are likely to make larger scale studies feasible in the near future.

## Supporting information

Supplemental Table 1 - Abundance Rankings

Supplemental Table 2 - Statistical Comparisons

Supplemental Table 3 - Fixation artefacts

## Abbreviations

ALDH1A1: aldehyde dehydrogenase 1 family member A1
ALDH2: aldehyde dehydrogenase 2
BCAM: basal cell adhesion molecule
BGN: biglycan
C3: Complement protein 3
C4A/B: complement proteins 4 A and B isoforms
CAA: 2-chloroacetamide
CD4: cluster of differentiation 4
CD36: cluster of differentiation 36
CD68: cluster of differentiation 68
CNN1: calponin 1
COL6A1: COL6A2, COL6A3, collagen VI-alpha1, -alpha2 and -alpha3 chains respectively
CTSB: cathepsin B
CVD: cardiovascular disease
DDA: data dependent acquisition
DDA-PASEF: data dependent acquisition with parallel acquisition-serial fragmentation
DIA: data independent acquisition
DIA-PASEF: data independent acquisition with parallel acquisition-serial fragmentation
DDM: *N*-dodecyl-β-D-maltoside
ECM: extracellular matrix
FFPE: formalin-fixed paraffin-embedded
FGA, FGB, FGG: fibrinogen alpha-, beta- and gamma- chains respectively
FLNA: filamin A
FTL: ferritin light chain
GCFHR: GTP cyclohydrolase 1 feedback regulatory protein
GO: gene ontology
GSEA: gene set enrichment analysis
HP: haptoglobin
HSPG2: heparin sulfate proteoglycan 2/perlecan
IGHM: immunoglobulin heavy constant mu
LAMB2: laminin beta-2 chain
LAMC1: laminin gamma-1 chain
LCM: laser-capture microdissection
LC-MS/MS: liquid chromatography-mass spectrometry
LDL: low-density lipoproteins
LUM: lumican
MARCO: macrophage receptor with collagenous structure
MMP12: matrix metalloproteinase 12
MYL6, MYL9: myosin light chains 6 and 9 respectively
MYH10, MYH11: myosin heavy chains 10 and 11 respectively
PASEF: parallel acquisition-serial fragmentation
PBS: phosphate-buffered saline
PCA: principal component analysis
POSTN: periostin
PRDX3: peroxiredoxin 3
PSMs: peptide spectral matches
TCEP: tris(2-carboxyethyl)phosphine
TEAB: tetraethyl ammonium bicarbonate
TFA: trifluoroacetic acid
TimsTOF: total ion mobility-time of flight
TNC: Tenascin C
VCAM1: vascular cell adhesion molecule 1
VCAN: versican
VWF: von Willebrand factor.

## Acknowledgements

This work was supported by grants from the Novo Nordisk Foundation (NNF13OC0004294 and NNF20SA0064214 to M.J.D.); the Lundbeck Foundation (R322-2019-2337 to L.F.G.) and the Novo Nordisk Foundation-University of Copenhagen BRIDGE scheme (to L.G.L.). The authors thank Max Sauerland for assistance with 3D printing of the Spin96 stage tip adapters.

## Data availability

The data supporting the findings of this study are available under controlled access due to ethical and legal restrictions related to the General Data Protection Regulation (GDPR). Qualified researchers may request access to the data through the University of Copenhagen. The data will be made available on the PRIDE database once an appropriate controlled access solution is implemented[47].

## Supplemental data

This article contains supplemental data.

## Supporting Data and Supplementary Methods

### Supplementary Methods

#### Tissue collection, embedding and sectioning

Carotid plaques were excised from patients as described recently [8]. Carotid artery atherosclerotic plaques from consecutive symptomatic patients, recently diagnosed with cerebral embolic ischaemia (stroke, transient ischaemic attack) or retinal ischaemia (complete vision loss, amaurosis fugax) were included. Patients were diagnosed, initially medically treated and subsequently referred for vascular surgery by departments of neurology. Preoperatively the patients received the recommended medical treatment, including dual antiplatelet therapy (acetylsalicylic acid 75 mg daily in combination with clopidogrel 75 mg daily), along with statins. Internal carotid artery endarterectomy (surgery) for all patients was carried out at the same department of vascular surgery. Plaques were removed *in toto* using standard surgical techniques and then immediately washed 3 times with cold PBS supplemented with 20 mM EDTA and 1X cOmpleteTM Protease Inhibitor Cocktail (Sigma-Aldrich #11836153001) and then placed in ice-cold deoxygenated (N_2_ gassed) PBS (prepared using Chelex-treated MilliQ water) containing antioxidants and preservatives (20 mM ethylenediaminetetraacetic acid (EDTA), 0.1 mM butylatedhydroxytoluene, 100 mM methionine, 1 mM azide and 1X cOmpleteTM Protease Inhibitor Cocktail).

Mouse heart tissue was obtained from B6.129P2-*Apoe^tm1Unc^* N11 male mice. Mice aged 8 weeks were purchased from Taconic Biosciences. Animals were housed in a temperature-controlled facility with a 12h light/dark cycle and received food and water *ad libitum*. The mice were euthanized at 24 weeks, perfused with cold saline before the heart was collected and prepared for cryosectioning as below.

Tissues were fixed in 4% formaldehyde in PBS overnight at room temperature followed by Tissue-Tek optimal cutting temperature medium (Sakura, 24 hrs, 4 °C). The tissue was then embedded in Tissue-Tek, flash frozen, and stored at -80°C. Polyethylene naphthalate membrane slides (1 mm, Zeiss) were treated with UV light for 1 h and coated with VECTABOND reagent (Vector Labs) according to manufacturer’s protocol. Plaques were cryosectioned at 10 µm thickness onto the coated membrane slides and stored at -80°C. Prior to staining, slides were dried either overnight at room temperature or at 37°C for 1 h to improve tissue adhesion. Tissue sections were stained with Mayer’s hematoxylin and eosin and air dried without mounting. Slides were the either used directly or stored at -80°C until use.

#### Laser capture microdissection

Laser capture microdissection and microscopy was performed with a Zeiss PalmRobo and 5x objective. Individual circular sections (diameter: 126, 178 or 252 μm) or square areas (225 x 225 µm) were cut with the laser and catapulted with a low energy diffuse laser pulse into 8 µL water drops pre-loaded into 8-well flat-cap PCR strips. Water droplets were loaded 8 to 16 wells at a time to avoid significant drying of the droplets during capture. For the mouse myocardial tissue, areas were obtained from up to 6 sequential sections. For the human plaques, morphologically matched square tissue areas were laser capture microdissected from up to 7 sequential sections.

Samples were processed as a ‘hanging-drop’ in a humid chamber (pipette box, lined top and bottom with wet paper towel) [20]. Briefly, flat 8-well PCR cap strips containing tissue captured in water droplets were placed face-up followed by addition of 100 mM triethylammonium bicarbonate (TEAB) and 0.05% *N*-Dodecyl-β-D-maltoside (DDM), returned face-down to the humid chamber and heated to 70°C for 40 min, placed face-up again, followed by reduction/alkylation with 10 mM tris(2-carboxyethyl)phosphine (TCEP) and 40 mM chloroacetamide (CAA) face-down by heating at 70°C for 20 min. 10 ng trypsin (modified sequencing grade, Promega) was added followed by overnight digestion face-down at 37°C, followed by acidification with 10% trifluoroacetic acid (TFA) and 96-well format stage tipping. Home-made stage-tips were prepared with C18 membranes (Affinisep AttractSPE) inserted into the bottom of P200 pipette tips [21]. 96-well stage-tipping was performed with transfer by centrifugation with a 3D-printed stage-tip adapter [48]. Samples were reconstituted in 5 µL of 5% formic acid (FA) in water.

#### Liquid-chromatography mass spectrometry proteomic analysis

Samples (1 µL loaded on-column) were analyzed on a TimsTOF Pro mass spectrometer (Bruker Daltonics) in the positive ion mode with a CaptiveSpray ion source on-line connected to a Dionex Ultimate 3000RSnano chromatography system (Thermo Fisher). Peptides were separated on an Aurora column (C18 1.6 μm, 5 cm, 150 μm ID, IonOpticks) at room temperature with a solvent gradient of 0.1% formic acid in water (Solvent A) and 99.9% acetonitrile/0.1% formic acid (Solvent B) over 15 min, at a flow rate of 1000 nL min^−1^ (total run time of 15 min). The mass spectrometer was operated in either data-dependent acquisition with parallel acquisition-serial fragmentation (DDA-PASEF) or data-independent acquisition parallel acquisition-serial fragmentation (DIA-PASEF) mode with 1.1 s cycle time and TIMS ramp time of 100 ms. MS scan range was set to 100–1700 *m*/*z* [26, 27]. DDA-acquired data was searched against the human UniProt reference proteome (UP000005640), including common contaminants, using Fragpipe version 18.0 (including MSFragger version 3.5 and Philosopher version 4.4.0), with the following settings: strict trypsin digestion (Trypsin/P) with 2 missed cleavages, precursor and fragment mass error ± 20 ppm, false discovery rate (FDR) 1% (protein and peptide level), cysteine carbamidomethylation (C +57.0215, fixed modification), *N*-terminal acetylation and methionine oxidation (*N*-term +42.0106 and M +15.9949, variable modifications) [28–33]. DIA-acquired data was searched against the human UniProt reference proteome (UP000005640) using DIA-NN version 1.9.1 with the following library-free settings: In silico digest at K/R, 1 missed cleavage, peptide length 7-35, charge 2-4, mass accuracy of 20/10 ppm (MS1/MS2), contaminants included, *N*-terminal methionine excision, cysteine carbamidomethylation (fixed modification), *N*-terminal acetylation and methionine oxidation (variable modifications)[49].

Fixation artefacts were assessed by MS-Fragger based open-search for peptide modifications in DDA data from 200.000 µm^2^ areas. Top modifications included formaldehyde adducts (delta m/*z* = + 12.00), formylation (delta m/*z* = + 27.99) and methylation (delta m/*z* = 14.02), and each represented less than 3 % of all peptide spectral matches (PSMs). These mass shifts are consistent with recent studies of bulk and laser captured fresh-fixed or FFPE tissue [50–52].

#### Statistical analysis and visualization

Data analysis and visualization of DIA data was performed using R (version 4.4.0) and R-Studio (Mac version 2024.04.1+748). Non-normalized precursor-level intensities were extracted from the DIA-NN report and analyzed using the MSqRobSum workflow to generate lists of differentially expressed proteins[53]. First, precursor intensities were log2-transformed, filtered (Q.Value < 0.01, PG.Q.Value < 0.01, contaminants from the cRAP database and keratins removed, precursors present in at least 5 out of 7 areas from sequential sections), vsn normalised and summarized into protein expression values with robust model-based summarization[54]. Peptides mapping to multiple proteins were summarized into groups of indistinguishable proteins. Finally, an analysis of differential protein expression between regions or sub-regions was carried out using a robust linear model approach. Proteins were considered as expressed differentially if the Benjamini-Hochberg adjusted p-value was < 0.01 [34].

Gene Set Enrichment Analysis (GSEA) was carried out using the clusterProfiler/ReactomePA packages with median-normalized log2 fold changes from the MSqRobSum analysis as input, and adjusted p-values <0.05 set as cut-off[55, 56].

## Supplementary Figures

**Supplementary Figure 1.**
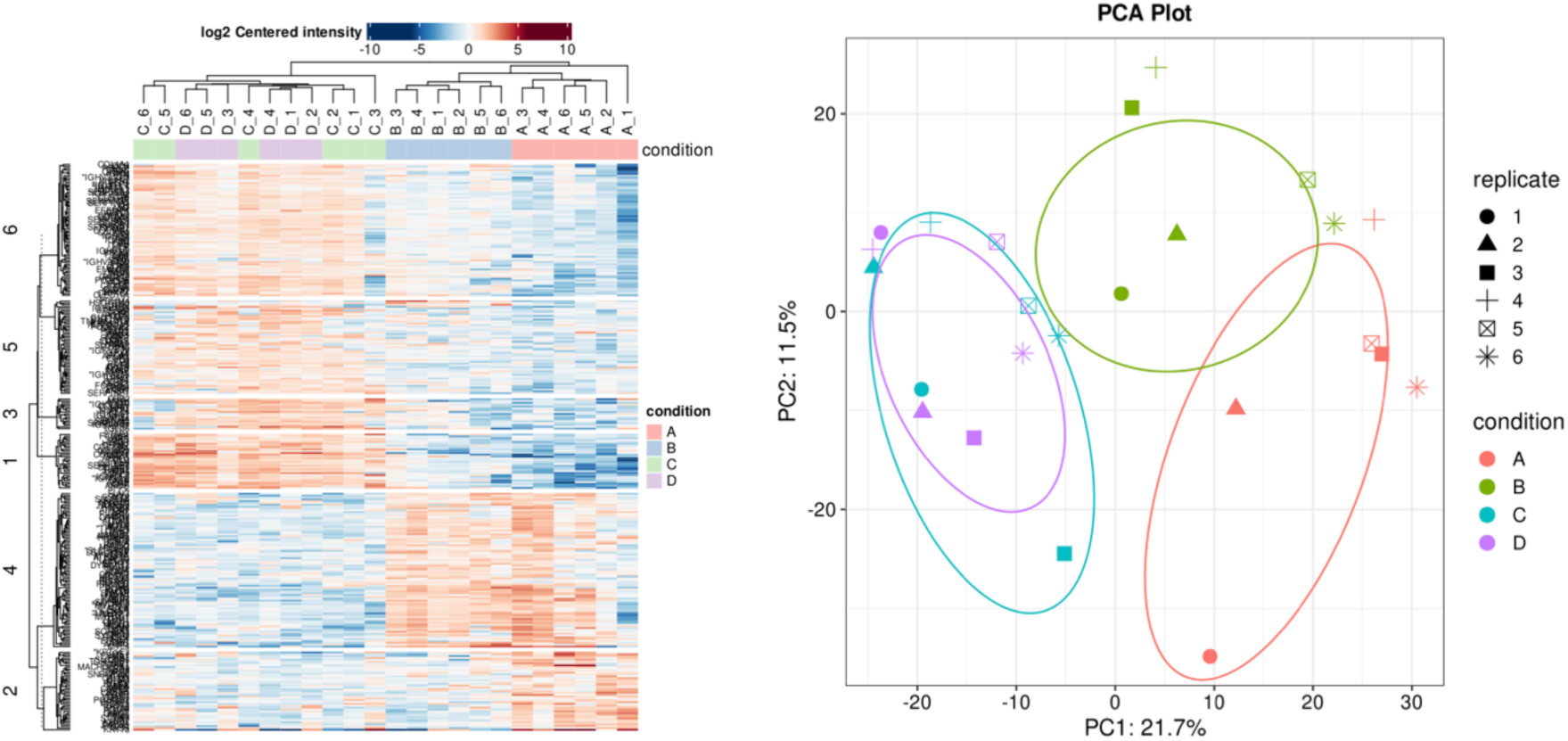
Heatmap and PCA of LCMS proteomic data from 4×2 array of human carotid artery plaque lumen to plaque gradient. 4×2 arrays were collected by LCM from sequential sections and samples were grouped based on distance from the lumen. A is from the luminal surface, D is the deepest into the plaque.

